# Effects of age, BMI and sex on the glial cell marker TSPO - a multicentre [^11^C]PBR28 HRRT PET study

**DOI:** 10.1101/564831

**Authors:** Jouni Tuisku, Pontus Plavén-Sigray, Edward C. Gaiser, Laura Airas, Haidar Al-Abdulrasul, Anna Brück, Richard E. Carson, Ming-Kai Chen, Kelly P. Cosgrove, Laura Ekblad, Irina Esterlis, Lars Farde, Anton Forsberg, Christer Halldin, Semi Helin, Eva Kosek, Mats Lekander, Noora Lindgren, Päivi Marjamäki, Eero Rissanen, Marcus Sucksdorff, Andrea Varrone, HRRT [^11^C]PBR28 study group, Juha Rinne, David Matuskey, Simon Cervenka, Members of HRRT [^11^C]PBR28 study group

## Abstract

**Purpose:** To investigate the effects of ageing, sex and body mass index (BMI) on translocator protein (TSPO) availability in healthy subjects using positron emission tomography (PET) and the radioligand [^11^C]PBR28.

**Methods:** [^11^C]PBR28 data from 140 healthy volunteers (72 males and 68 females; n=78 with HAB and n=62 MAB genotype; age range 19-80 years; BMI range 17.6 - 36.9) were acquired with High Resolution Research Tomograph at three centres: Karolinska Institutet (n=53), Turku PET centre (n=62) and Yale University PET Center (n=25). The total volume of distribution (V_T_) was estimated in global grey matter, frontal, temporal, occipital and parietal cortices, hippocampus and thalamus using multilinear analysis 1. The effects of age, BMI and sex on TSPO availability were investigated using linear mixed effects model, with TSPO genotype and PET centre specified as random intercepts.

**Results:** There were significant positive correlations between age and V_T_ in the frontal and temporal cortex. BMI showed a significant negative correlation with V_T_ in all regions. Additionally, significant differences between males and females were observed in all regions, with females showing higher V_T_. A subgroup analysis revealed a positive correlation between V_T_ and age in all regions in male subjects, whereas age showed no effect on TSPO levels in female subjects.

**Conclusion:** These findings provide evidence that individual biological properties may contribute significantly to the high variation shown in TSPO binding estimates, and suggest that age, BMI and sex can be confounding factors in clinical studies.

## Introduction

The translocator protein (TSPO) is an 18 kDa protein structure located on the outer mitochondrial membrane. In the brain, TSPO is expressed in microglia, astrocytes, endothelial and smooth muscle cells and even neurons (although in lower levels compared to tissues which are essential for steroid production or lipid storage and metabolism) [1, 2]. Although TSPO is present throughout the brain under normal physiological conditions [3–5], *in vitro* studies have shown that TSPO expression increases in response to immune activation [6]. TSPO radioligands have therefore been used together with molecular imaging techniques to study brain immune activation.

A limitation of clinical TSPO PET studies is that TSPO shows high interindividual variability, even after accounting for the effect of the TSPO gene polymorphism [7, 8]. It is not fully known to what extent this variability is influenced by individual physiological properties, such as age, body mass index (BMI) and sex. TSPO is involved in the transport of cholesterol across the mitochondrial membrane, a requirement for steroid synthesis; additionally it is involved in immunomodulation as well as mitochondrial respiration and metabolism [9]. These processes are likely to be related to ageing, as well as hormonal steroid function [10]. However, previous *in vivo* human research using positron emission tomography (PET) imaging to investigate influences of age and sex on TSPO expression have been inconclusive. Studies have showed both higher TSPO levels with increasing age [11–14], and no effect [15–17]. Similarly, no effects of sex or BMI on TSPO expression were found in a recent TSPO study of healthy control subjects [13], whereas higher TSPO levels were observed in females in a clinical study with small sample size [18].

*In vivo* PET studies usually involve small sample sizes, and this low statistical power adds difficulty to detect significant effects. In order to more conclusively examine the effects of ageing, BMI and sex on TSPO availability in healthy individuals, we combined TSPO radioligand [^11^C]PBR28 data acquired using a High Resolution Research Tomograph (HRRT) from three different institutes. This resulted in the largest TSPO PET sample to date of healthy control subjects imaged using the same radioligand and PET system.

## Materials & Methods

[^11^C]PBR28 data from 140 healthy volunteers were enrolled at three centres: Karolinska Institutet (KI), n = 53, Turku PET Centre (TPC), n = 62 and Yale University PET Center (Yale), n = 25. Descriptive statistics for age, BMI, sex and TSPO genotype for each centre are presented in Table 1.

**Table 1.**
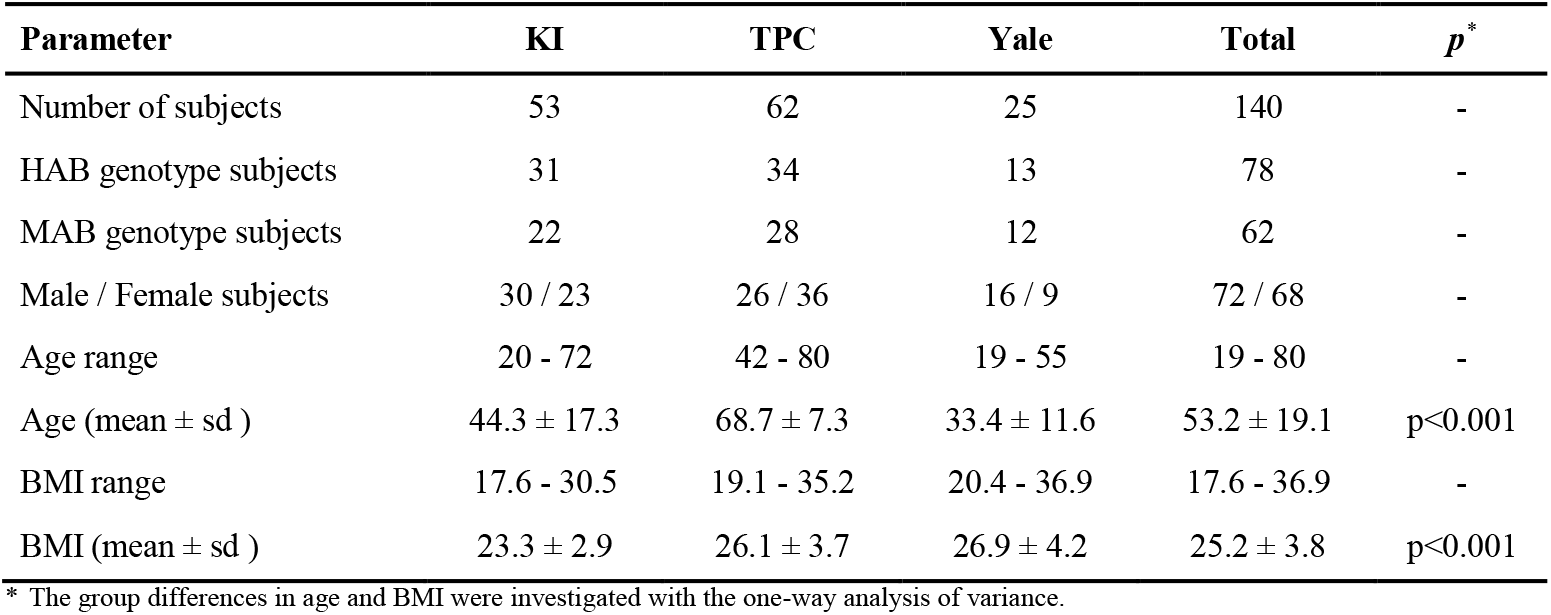
Study subject characteristics of Karolinska Institutet (KI), Turku PET Centre (TPC) and Yale University PET Center (Yale) data.

| Parameter | KI | TPC | Yale | Total | *p*<sup>*</sup> |
| --- | --- | --- | --- | --- | --- |
| Number of subjects | 53 | 62 | 25 | 140 | - |
| HAB genotype subjects | 31 | 34 | 13 | 78 | - |
| MAB genotype subjects | 22 | 28 | 12 | 62 | - |
| Male / Female subjects | 30 / 23 | 26 / 36 | 16 / 9 | 72 / 68 | - |
| Age range | 20 - 72 | 42 - 80 | 19 - 55 | 19 - 80 | - |
| Age (mean ± sd ) | 44.3 ± 17.3 | 68.7 ± 7.3 | 33.4 ± 11.6 | 53.2 ± 19.1 | *p*<0.001 |
| BMI range | 17.6 - 30.5 | 19.1 - 35.2 | 20.4 - 36.9 | 17.6 - 36.9 | - |
| BMI (mean ± sd ) | 23.3 ± 2.9 | 26.1 ± 3.7 | 26.9 ± 4.2 | 25.2 ± 3.8 | *p*<0.001 |

\* The group differences in age and BMI were investigated with the one-way analysis of variance.

Exclusion criteria for participation at all centres were evidence of a current or previous major psychiatric illness (e.g., schizophrenia or major depression), alcohol or drug dependence, or history of a serious somatic or neurological illness. All subjects had a physical exam and medical history to confirm criteria. Subjects who had medication for diabetes mellitus (n=7, exclusively in Turku data) were excluded from the initial study sample of 147 individuals. Three subjects of Turku data were smoking regularly, and for several Turku data subjects the current smoking status was unknown (n=10). A subset of Turku data subjects also used cholesterol-lowering statins (n=11) and hormonal replacement medication (n=2). Prior to imaging, all participants were genotyped for the rs6971 polymorphism of the TSPO gene and categorized as mixed-affinity binders (MABs) or high-affinity binders (HABs) utilizing methods described previously [7, 19]. Low-affinity binders were excluded from this study.

Structural MR imaging (MRI) was performed to exclude individuals with anatomical abnormalities and to allow for the delineation of anatomical regions for the PET data analysis after coregistration. T1-weighted MRI scans were acquired using a 3T scanner (Discovery MR750 system (GE, Milwaukee, WI, at KI; Philips Ingenuity TF PET/MR, Philips Medical Systems, Cleveland, OH, USA, at TPC; Trio system, Siemens Medical Solutions, Malvern, Pennsylvania, at Yale).

All PET examinations were performed on a HRRT scanner (Siemens/CTI, Knoxville, TN, USA). The [^11^C]PBR28 synthesis in TPC is described in the online resource. In KI and Yale the [^11^C]PBR28 was prepared as previously described [19, 20]. The injection dose was administered as a rapid intravenous bolus, where the mean (sd) of injected activity and molar activity at the time of injection, respectively, at each centre was 413.5 (53.4) MBq and 328.5 (148.0) MBq/nmol at KI, 494.3 (18.9) MBq and 286.9 (120.0) MBq/nmol at TPC and 575.3 (151.6) MBq and 178.7 (164.0) MBq/nmol at Yale. Data were acquired while participants were at rest over 70, 75 or 90 minutes, respectively at TPC, KI and Yale. The present analysis was restricted to 70 minutes at TPC, and 75 minutes at KI and Yale. The further details of PET data acquisition, reconstruction and frame lengths, as well as arterial blood sampling are presented in the online resource.

All 4D PET data were preprocessed in a similar manner, in which PET images were realigned and coregistered to anatomical MR images using SPM12 software (Wellcome Trust Centre for Neuroimaging, London, UK) running in MATLAB (The Mathworks, Natick, MA), or Bioimage Suite (version 2.5; http://www.bioimagesuite.com). The MR images were further segmented into tissue classes using SPM. The grey matter segment was thresholded (GM>0.5) and then used as the main region of interest for the statistical analysis. Additionally, six automated ROIs (frontal, temporal, occipital and parietal cortices, hippocampus and thalamus) were generated by using the Automated Anatomical Labeling (AAL) template [21]. These AAL ROIs were first coregistered to summed PET images via the deformation fields obtained from the MRI segmentation and then multiplied with the thresholded GM segment, before the ROI extraction.

The total volume of distribution (V_T_) was estimated using multilinear analysis 1 [22] (t* = 30min) with the metabolite and the delay corrected arterial plasma curves as an input function. Uniform weights were used in the MA1 analysis at KI and Turku, whereas at Yale the weights were based on the noise equivalent counts in each frame. The normality of the regional V_T_ estimates were inspected with Shapiro-Wilk test of normality and the Q-Q plot. The main research question of this study was to examine the associations between [^11^C]PBR28 binding and age, sex as well as BMI. This was done using a linear mixed effects model, with TSPO genotype and PET centre specified as random effects, allowing their intercepts to vary freely. The results from this model are reported as “confirmatory”, as the associations were hypothesized *a priori*. Having observed the results from the confirmatory analysis, we performed a set of follow-up tests, examining males and females separately, as well as sex-age interaction and centre effects. The results of these analyses are reported as “exploratory”, as they are all *post-hoc*. The results are presented as unstandardized regression effect estimates with 95% confidence intervals. All statistical analyses were conducted using lme4 and lmerTest packages in R (version 3.5.0 “Joy in Playing”). Previous studies have shown high correlations between [^11^C]PBR28 V_T_ estimates in different ROIs [23] and in this dataset all inter-regional Pearson’s correlations were >0.96 (all p<10^−15^). Hence, we considered the ROI analyses to be highly dependent comparisons and set the alpha threshold to 0.05 (two-tailed).

## Results

There were significant differences in the age and BMI between centres (Table 1), but there were no significant differences in these variables between HABs and MABs, or between males and females (Table 2). Figure 1 illustrates the relationships of age, BMI and sex to grey matter [^11^C]PBR28 V_T_. The Shapiro-Wilk test and the Q-Q plot indicated that the residuals of the statistical model were not normally distributed (see Online Resource). Hence, all V_T_ values were log-transformed prior to being inserted into the linear-mixed effect model, which then fulfilled the assumption of normally distributed residuals. Below, the results from log-transformed V_T_ values are presented.

**Fig. 1.**
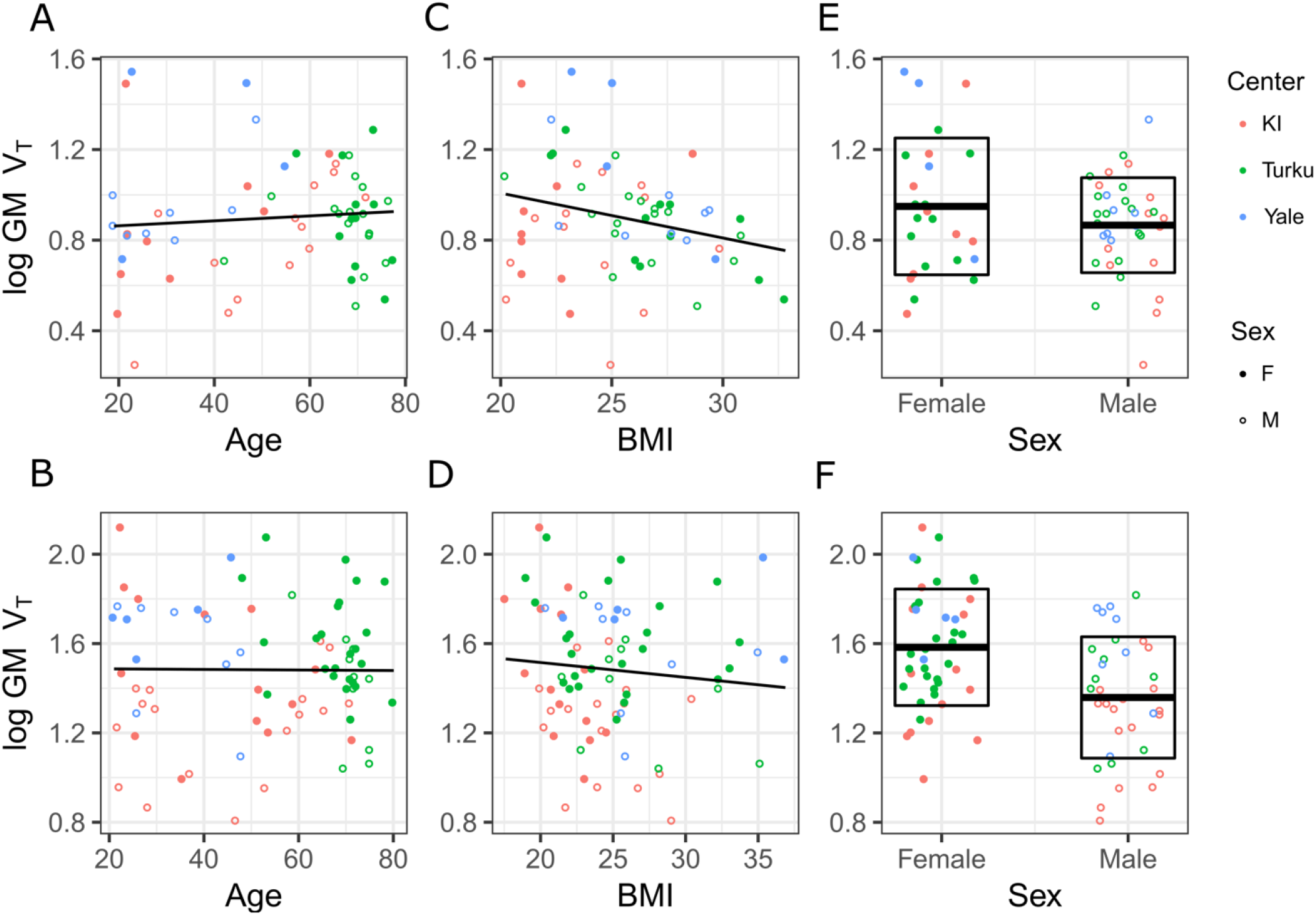
The relationships of age, BMI and sex to log-transformed grey matter [^11^C]PBR28 V_T_. Age vs. [^11^C]PBR28 GM V_T_ in MAB genotype (**A**) and HAB genotype (**B**). BMI vs. [^11^C]PBR28 GM V_T_ in MAB genotype subjects (**C**) and HAB genotype subjects (**D**). Sex vs. [^11^C]PBR28 GM V_T_ in MAB genotype subjects (**E**) and HAB genotype subjects (**F**). Box-plots in (**E**) and (**F**) represent the data means and standard deviations for both sexes. Using the raw, untransformed V_T_ values the results were similar to those presented above, except for the V_T_-age association that was not statistically significant (see online resource).

**Table 2.** Study sample characteristics grouped by TSPO genotype (**A**) and sex (**B**).

| (A) | Demographic | HAB | MAB | $p^*$ | (B) | Female | Male | $p^*$ |
| --- | --- | --- | --- | --- | --- | --- | --- | --- |
|  | n | 78 | 62 | - |  | 68 | 72 | - |
|  | n (Males/Females) | 35 / 43 | 37 / 25 | - |  | - | - | - |
| | Age (mean ± sd) | 53.2 ± 18.9 | 53.1 ± 19.6 | 0.969 | | 54.0 ± 19.6 | 52.4 ± 18.8 | 0.632 |
| | BMI (mean ± sd) | 24.9 ± 4.2 | 25.5 ± 3.1 | 0.394 | | 24.7 (4.3) | 25.6 ± 3.3 | 0.195 |

\*The group differences in age were investigated with the Student's t-test.

### Confirmatory results

The regional fixed effect estimates with confidence intervals and p-values for the association between [^11^C]PBR28 V_T_ and age, BMI and sex are presented in Table 3. There was a significant positive association between age and log-transformed V_T_ in frontal and temporal cortex, but not in any other regions. Higher BMI levels were associated with lower log-transformed V_T_ in all regions. Additionally, significant differences between males and females were observed in all regions, with females showing higher [^11^C]PBR28 binding. For example, an increase of one year of age predicted a 0.4% increase (100*(e^0004^ − 1) = 0.4) in temporal cortex V_T_. Correspondingly, an increase of one BMI unit predicted a 2.4% decrease (100*(e^0024^ − 1) = −2.4) in temporal cortex V_T_. For females, temporal cortex V_T_ was 16.3% higher (100*(e^0151^ − 1) = 16.3) compared to males.

**Table 3.** Fixed effects estimates with 95% confidence intervals and p-values for age, BMI and sex as regressors for regional [^11^C]PBR28 log-transformed V_T_. N = 140 for all analyses.

| Region | Age ($\beta$, [CI], p) | BMI ($\beta$, [CI], p) | Sex ($\beta$, [CI], p)* |
| --- | --- | --- | --- |
| **Gray matter** | 0.003, [-0.001,0.007], p=0.080 | -0.022, [-0.034,-0.010], p<0.001 | 0.155, [0.075,0.235], p<0.001 |
| **Frontal cortex** | 0.004, [0.000,0.008], p=0.034 | -0.021, [-0.033,-0.009], p=0.001 | 0.164, [0.082,0.246], p<0.001 |
| **Temporal cortex** | 0.004, [0.000,0.008], p=0.030 | -0.024, [-0.036,-0.012], p<0.001 | 0.151, [0.069,0.233], p<0.001 |
| **Occipital cortex** | 0.002, [-0.002,0.006], p=0.167 | -0.019, [-0.031,-0.007], p=0.001 | 0.161, [0.081,0.241], p<0.001 |
| **Parietal cortex** | 0.003, [-0.001,0.007], p=0.086 | -0.017, [-0.029,-0.005], p=0.004 | 0.172, [0.092,0.252], p<0.001 |
| **Hippocampus** | 0.002, [-0.002,0.006], p=0.349 | -0.026, [-0.038,-0.014], p<0.001 | 0.140, [0.054,0.226], p=0.002 |
| **Thalamus** | 0.003, [-0.001,0.007], p=0.067 | -0.023, [-0.035,-0.011], p<0.001 | 0.139, [0.051,0.227], p=0.002 |

\*Coding: Males=0, Females=1

### Exploratory results

Due to the sex differences in V_T_, the data were divided into two subgroups for studying the effects of age and BMI separately for both sexes. This subgroup analysis showed a significant positive association between age and log-transformed V_T_ in all regions in male subjects, but not in female subjects (Table 4). There was also a significant negative association between BMI and [^11^C]PBR28 binding in males in all regions, but for females the association was significant only in the temporal cortex, hippocampus and thalamus. For example, an increase of one year of age predicted a 0.7% and a 0.1% increase in temporal cortex V_T_, respectively for males and females. Correspondingly, an increase of one BMI unit predicted 2.6% and 1.9% decrease in temporal cortex V_T_, respectively for males and females.

**Table 4.**
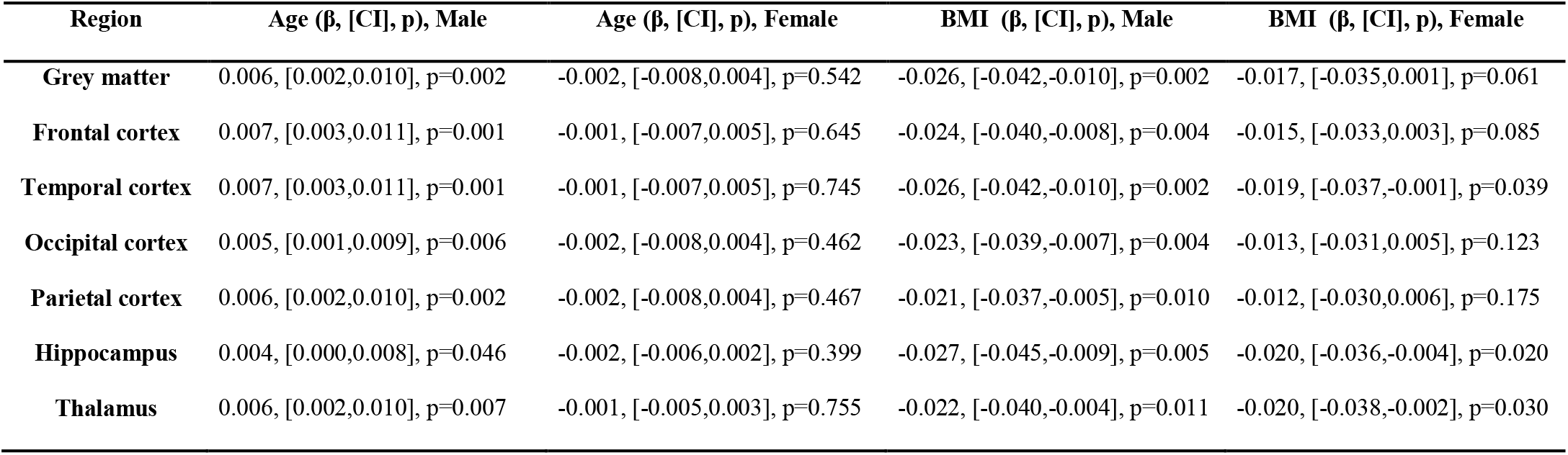
Fixed effects estimates with 95% confidence intervals and p-values for age and BMI as regressors for regional [^11^C]PBR28 log-transformed V_T_ in a separate subgroup analysis for males (N = 72) and females (N = 68).

| Region | Age ($\beta$, [CI], p), Male | Age ($\beta$, [CI], p), Female | BMI ($\beta$, [CI], p), Male | BMI ($\beta$, [CI], p), Female |
| --- | --- | --- | --- | --- |
| **Grey matter** | 0.006, [0.002,0.010], p=0.002 | -0.002, [-0.008,0.004], p=0.542 | -0.026, [-0.042,-0.010], p=0.002 | -0.017, [-0.035,0.001], p=0.061 |
| **Frontal cortex** | 0.007, [0.003,0.011], p=0.001 | -0.001, [-0.007,0.005], p=0.645 | -0.024, [-0.040,-0.008], p=0.004 | -0.015, [-0.033,0.003], p=0.085 |
| **Temporal cortex** | 0.007, [0.003,0.011], p=0.001 | -0.001, [-0.007,0.005], p=0.745 | -0.026, [-0.042,-0.010], p=0.002 | -0.019, [-0.037,-0.001], p=0.039 |
| **Occipital cortex** | 0.005, [0.001,0.009], p=0.006 | -0.002, [-0.008,0.004], p=0.462 | -0.023, [-0.039,-0.007], p=0.004 | -0.013, [-0.031,0.005], p=0.123 |
| **Parietal cortex** | 0.006, [0.002,0.010], p=0.002 | -0.002, [-0.008,0.004], p=0.467 | -0.021, [-0.037,-0.005], p=0.010 | -0.012, [-0.030,0.006], p=0.175 |
| **Hippocampus** | 0.004, [0.000,0.008], p=0.046 | -0.002, [-0.006,0.002], p=0.399 | -0.027, [-0.045,-0.009], p=0.005 | -0.020, [-0.036,-0.004], p=0.020 |
| **Thalamus** | 0.006, [0.002,0.010], p=0.007 | -0.001, [-0.005,0.003], p=0.755 | -0.022, [-0.040,-0.004], p=0.011 | -0.020, [-0.038,-0.002], p=0.030 |

The interaction between age and sex was confirmed by adding an interaction term to the main effects model (fixed effects estimates are presented in the online resource). This analysis revealed a significant age and sex interaction in all regions except the occipital cortex and hippocampus. The fixed effect estimates of the association between age and log-transformed V_T_ in the grey matter for males and females are illustrated in Figure 2.

**Fig. 2.**
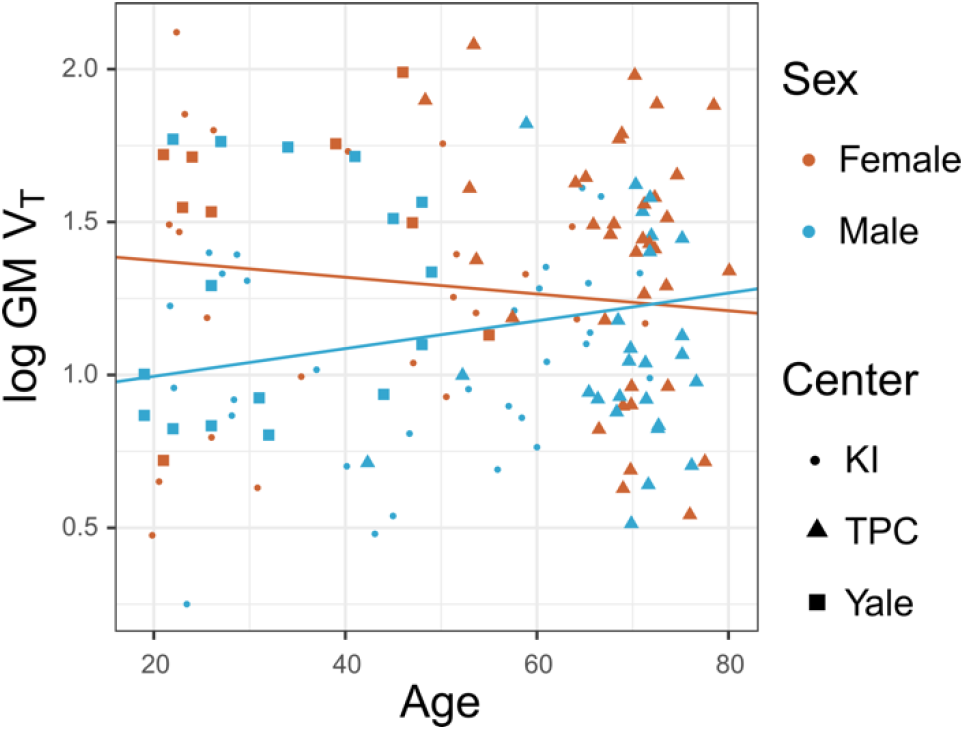
The relationship between [^11^C]PBR28 log-transformed V_T_ and age for males and females in the grey matter. The regression lines denote the fixed effects from two linear mixed effects models including either females or males, with TSPO genotype and PET center specified as random intercepts.

### Robustness test

To examine the robustness of our results, we performed the main analysis also by excluding the subjects who had cholesterol medication (n=11), hormone replacement therapy (n=2) and who were current smokers (n=3) (presented in the online resource; 16 excluded subjects, thus resulting in a total number of subjects of 124). The results were similar compared to the main analysis without exclusions.

Because of the differences in age and BMI distributions between centres, it can’t be excluded that the overall age or BMI effects are attributed to only one centre. Therefore, a possible centre effect was studied in grey matter by inserting centre as a dummy predictor and as an interaction-effect to age, BMI and sex. The results were comparable to the main analysis. The age interaction estimate for Turku data was statistically significant, whereas the other interaction estimates were not significant (the results are presented in the online resource).

## Discussion

In this multicentre collaboration study, constituting the largest harmonized TSPO PET sample to date, the results indicate that higher [^11^C]PBR28 V_T_ is associated with lower BMI. Higher V_T_ was found in females, as compared to males. Higher age was significantly associated to higher V_T_ in frontal and temporal cortices, partly replicating a previous [^11^C]PBR28 study [13]. A post hoc analysis however revealed that this association was present only in male subjects. These findings provide evidence that individual biological factors contribute significantly to the high variability seen in V_T_ estimates, and suggest that age, BMI and sex can be confounding factors in clinical designs.

It has been suggested that with increasing age, microglia activate slower and exhibit less “surveying” morphology [24]. Several post-mortem human studies have indicated age-related alterations in microglia morphology and function as well as age-associated increases in total number of activated pro-inflammatory microglia phenotype [25–28]. Ageing has also been shown to be associated with increases in peripheral markers of inflammation [29–32]. In contrast, previous PET studies on brain TSPO in relation to age have been inconclusive [11, 13–15, 17, 33, 34]. However, the sample sizes have generally been small, limiting the conclusions that can be drawn. By using a several-fold larger dataset, a second-generation TSPO radioligand and full arterial sampling, our observation of a positive effect of age on [^11^C]PBR28 binding in the frontal and temporal cortices is congruent with some of the previous studies, where positive correlations between age and TSPO binding were found in the thalamus [11], whole cortical GM [14] and in several cortical and subcortical areas [13, 33]. Importantly, the present results were driven solely by the males, which could explain the conflicting results of no observed age effect found in other studies [15, 17, 34], where mixed samples of males and females were included.

Previous preclinical studies have shown also sex-related differences in microglial function [35, 36]. These findings are supported by our observation of females showing higher [^11^C]PBR28 V_T_ compared to males. Also, the potential link between TSPO and steroidogenesis could partly explain the association between immune function and sex steroid hormones [37]. Particularly, glial cells express receptors for estrogens and androgens, suggesting that there is an interplay between sex steroid hormones and the neuroinflammatory response [38–40]. As a consequence, an age-related change in both androgens and female sex hormones could explain the observed interaction of age and sex in [^11^C]PBR28 binding, where male TSPO levels increase with age, eventually reaching the higher TSPO level of females. Our finding of age-related sex difference in [^11^C]PBR28 V_T_ may also provide a clue to the observed sex-specific differences of neuroinflammatory autoimmune diseases such as multiple sclerosis, which are more prevalent in younger females than in males [41].

While the immunostaining assays of a preclinical model of obesity have suggested that high-fat diet induced obesity increases neuroinflammation in the hypothalamus and hippocampus [42–45], in contrast we observed decreased ^11^C]PBR28 V_T_ with higher BMI. A possible explanation for this could be that the peripheral TSPO expression is dysregulated in obese subjects, and that this effect is reflected also in the brain. Interestingly, a recent *in vitro* study on adipocytes, which are considered to be a major regulatory cell in obesity, suggested that TSPO expression is essential for the maintenance of the healthy adipocyte functions, and that TSPO activation in adipocytes improves their metabolic status in regulating glucose homeostasis [46]. On the other hand, a dysregulation of TSPO expression has been reported previously in a preclinical study, where the total number of available TSPO binding sites (Bmax) was significantly decreased in brown adipose tissue cells of obese mice compared to the lean controls, whereas there was no difference in the affinity of the ligand (Kd) [47]. Although our results support an involvement of TSPO in obesity, it is currently unknown if this effect is related to TSPO expression in microglia, astrocytes, endothelial or even adipose cells.

Another explanation for the observed negative association between TSPO and BMI could be related to the increased level of endogenous ligands such as cholesterol and porphyrins [48], for which TSPO has high affinity. Obese subjects often have higher levels of serum cholesterol [49], which could induce competition between TSPO ligand and cholesterol between available TSPO binding sites. This hypothesis is supported by a previous study, where higher plasma cholesterol levels were shown to be associated with lower whole brain [^11^C]PBR28 V_T_ in healthy control subjects and alcohol use disorder patients [50]. Cholesterol medication such as statins are expected to affect serum cholesterol level. However, excluding these individuals (n=11) from the analysis did not change our results. Taken together, further studies on the role of TSPO and neuroinflammation in obesity are warranted to address these questions.

Our study includes some limitations. Although the subjects are imaged using the same PET scanner with similar scanning protocols, the data are acquired in different institutes with significantly different age and BMI distributions between the study subjects. In addition, the scanner hardware and software differences might increase the data variability and even influence the results on TSPO availability-associations. However, because of the nested structure of the data, we strived to control for the possible centre differences statistically by modelling a random intercept for each centre. Additionally, although all subjects were categorized as healthy controls, there are several confounding factors, which might contribute to the TSPO availability. Several older subjects were using statins to control cholesterol levels, which could reduce the competition between the TSPO radioligand and cholesterol and would improve the radioligand availability in the brain. Also, smokers have been reported to have decreased TSPO availability [51], which could contribute also to our results, as the smoking status of several subjects at the time of PET acquisition was unknown. Importantly, the main analysis results remained the same after excluding the subjects who were using cholesterol lowering or hormonal replacement medication, or whose current smoking status was positive. Thus, it is less likely that these subjects have significant effects on our age, BMI and sex findings.

In summary, we show a positive effect of ageing on TSPO, partly replicating a previous study using [^11^C]PBR28. In addition, we found that TSPO levels were associated to sex and BMI, which have not been demonstrated previously. In addition to identifying important confounding factors for clinical TSPO studies, the present study shows how a multicentre collaboration can mitigate the problem of small sample sizes in PET research, providing increased power to detect clinically relevant effects. The large cohort of healthy control subjects provides an excellent pool to be used as a reference in future studies applying [^11^C]PBR28 for evaluation of various brain disorders.

## Compliance with ethical standards

### Ethical approval

All procedures performed in studies involving human participants were in accordance with the ethical standards of the institutional and/or national research committee and with the principles of the 1964 Declaration of Helsinki and its later amendments or comparable ethical standards.

### Informed consent

Informed consent was obtained from all individual participants included in the study.

### Funding

PET studies at KI were funded by grants from the Swedish Science Council. PET studies at Turku PET Centre were funded by grants from Sigrid Juselius foundation, Pro Humanitate foundation, Finnish Cultural Foundation, Academy of Finland (#310962), Governmental Clinical Grants (VTR) and The Finnish Parkinson Foundation. PET studies at Yale were funded by grants from Nancy Taylor Foundation and SNMMI molecular imaging research grant for junior faculty. JT is supported by the Alfred Kordelin foundation, the Instrumentarium science foundation, the Orion research foundation, the Paulo foundation, the Päivikki and Sakari Sohlberg foundation and Turku University Hospital Foundation. SC is supported by Swedish Research Council (Grant No. 523-2014-3467). MKC is supported by Eli Lilly and company grant. NL was supported by the Finnish Cultural Foundation, Yrjö Jahnsson Foundation, Turku University Foundation, and Finnish Brain Foundation. KC is supported by U.S. Department of Veterans Affairs National center for posttraumatic stress disorder. ML is supported by Hedlund Foundation, Swedish Heart-Lung Foundation, Swedish Asthma and Allergy Association and the Swedish Research Council. HA is supported by the Finnish Parkinson Foundation. LA is supported by the Academy of Finland and the Sigrid Juselius foundation.

### Availability of data and material

The datasets used and analysed during the current study may be available from the principal investigators on reasonable request.

### Authors’ contributions

DM, JR, SC supervised and designed the study; PSS designed the statistical model; JT and PPS performed the statistical analysis of all data; BP contributed to the statistical analysis of the data; AF and PPS performed the KI data processing at KI; JT performed the Turku data processing; EK and ML performed the subject recruitment for PET scanning at KI; LF, and AV participated in the generation and analysis of the [11C]PBR28 data at KI; LA, HAA, AB, LE, NL and MS performed the subject recruitment and PET scanning at Turku; REC, MKC, KPC, IE, JDG, AH, YH and CMS participated in generation and analysis of the [11C]PBR28 data at Yale; CH performed the radiochemistry at KI; SH performed the radiochemistry at Turku; NN supervised Yale PET radiochemistry; PM performed the metabolite analysis for Turku data; JT, DM, JR, PPS and SC drafted the manuscript; ECG, LA, LE, SH, ML, ER participated in manuscript writing; All authors interpreted the results, critically revised the article, and approved of the final version for publication.

## Acknowledgements

We would like to thank all study participants and the staff at the Karolinska Institutet PET Centre, Turku PET Centre and Yale University PET Center for their assistance. We thank also Tomi Karjalainen for the help in the analysis of Turku data.

