## Supplementary material for "Effects of age, BMI and sex on the glial cell marker TSPO - a multicentre [^11^C]PBR28 HRRT PET study"

[9]*Members of HRRT [$^{11}$C]PBR28 study group are listed in the Acknowledgments before References.*

** Denotes shared last author contributions*.

**Corresponding author:**

Jouni Tuisku,

**The details of [$^{11}$C]PBR28 synthesis in Turku PET Centre**

In TPC the [$^{11}$C]PBR28 was produced using an adaption of a published method [1]. Cyclotron-produced [$^{11}$C]methane was halogenated by gas phase reaction into [$^{11}$C]methyl iodide, converted online into [$^{11}$C]methyl triflate and bubbled in a solution of desmethyl precursor (0.30-0.40 mg, PharmaSynth AS, Tartu, Estonia) in acetone (200 µl) and sodium hydroxide (0.5 M NaOH, 2.5 µl) at 0 °C. After 3 minutes reaction at 80 °C, the reaction product was purified using a semi-preparative high performance liquid chromatography (HPLC) column (ACE C18-HL, 5 µm, 10 × 250 mm, Advanced Chromatography Technologies Ltd, Aberdeen, Scotland) and $CH_3CN$ in 0.02 M $KH_2PO_4$ (43:57) mobile phase at 7 ml/min flow rate. The eluent was monitored for 254 nm UV absorbance and radioactivity. Collected product fraction was diluted and loaded into a solid phase extraction (SPE) C18 cartridge (Sep-Pak® Light, Waters Corp., Milford, MA) and the cartridge was washed. Dilution and washing were done using five percent ethanol in water solution. The product was extracted with ethanol from the SPE cartridge, diluted with 0.1 M phosphate buffer solution into < 10% ethanol level and finally sterile filtered (Millex GV, 0.22 µm polyvinylidene fluoride membrane, 33 mm, Merck Millipore). Determination of identity, radiochemical purity, and mass concentration was performed using analytical HPLC column (Kinetex XB-C18, 2.6 µm, 3.00 × 50 mm, Phenomenex Inc., USA ), $CH_3CN$ in 0.02 M $KH_2PO_4$ (35:65) mobile phase, 1 ml/min flow rate, 5 min run time and detectors in series for UV absorption (210 nm) and radioactivity. Radiochemical purities were better than 99.9% and mean (sd) molar activity at the end of synthesis was 600 (220) MBq/nmol.

**The details of PET data acquisition, reconstruction and arterial blood sampling**

All PET examinations were performed on a HRRT scanner (Siemens/CTI, Knoxville, TN, USA), which acquired 207 slices separated by 1.28 mm with a reconstructed image resolution of approximately 2.5 mm [2]. A transmission scan using $^{137}Cs$ point source was obtained for attenuation correction of the PET emission data prior to tracer injection. At KI and TPC, a thermoplastic mask was used with a head fixation system to minimize subject movement artefacts. At Yale, an optical motion detector (Vicra, NDI Systems, Waterloo, Ontario, Canada) was utilized for motion correction during image reconstruction. List–mode data were reconstructed using 3D Ordinary Poisson Ordered Subset Estimation Maximization with 10 iterations and 16 subsets at KI, 8 iterations and 16 subsets at TPC, and 2 iterations and 30 subsets at Yale. At KI and Yale the reconstruction included additional modeling of the point-spread function [3, 4], and at Yale the subject motion was corrected within reconstruction employing the MOLAR algorithm [4]. The present analysis was restricted to 70 minutes at TPC, and 75 minutes at KI and Yale, with frame lengths of 9 x 10s, 2 x 15 s, 3 x 20 s, 4 x 30 s, 4 x 60 s, 4 x 180 s and 9 x 360 s at KI; 6 x 30 s, 3 x 60 s, 2 x 120s and 12 x 300s at TPC and 6 x 30 s, 3 x 60 s, 2 x 120s and 13 x 300s at Yale.

Arterial blood sampling were carried out as described previously [5, 6] in order to obtain an arterial input function. In short, an automated blood sampling system (ABSS, Allogg AB, Mariefred, Sweden) was used during the first 5 min of each PET measurement. Arterial blood samples were also drawn manually at 1, 3, 5, 7, 9, 10.5, 20, 30, 40, 50, 60, 70 min at KI; at 4, 6, 8,10, 15, 20, 25, 30, 40, 50 and 70 at TPC, and at 8, 12, 15, 20, 25, 30, 40, 50, 60, 75 at Yale, to allow for radioactivity sampling and metabolite analysis. A metabolite corrected arterial input curve was created by first merging the ABSS blood curve with the manually drawn blood samples. The blood samples were centrifuged, and radioactivity in plasma was measured. Following this, a plasma-to-blood ratio curve was established and multiplied with the whole blood curve, to create the plasma curve over the entire examination. At KI and Yale, individual plasma-to-blood ratio data was used in the conversion, whereas at TPC, Hill type population-based plasma-to-blood ratio curves corresponding to both genotypes were utilized. Next, for all centers a Hill type function was fitted to each individual's parent fraction measurements, after which the metabolite corrected plasma time-activity curves were calculated by multiplying the uncorrected plasma curves with the estimated model curves. The differences in appearance times of radioactivity between PET and arterial plasma time-activity curves (TACs) were corrected by first estimating the delay of the arterial plasma TAC, which produced the best fit of two tissue compartment model to whole brain TAC and then shifting the arterial plasma TAC accordingly.

**Test of statistical assumptions and raw $V_T$ values**

For the linear-mixed-effects models (see Materials and Methods in the main text), the ensuing residuals were not normally distributed, as indicated by the Shapiro-Wilkow test (sTable 1) and the Q-Q plot (sFigure 1 for the grey matter $V_T$). For this reason, $V_T$ values for all regions were log-transformed and the statistical analyses were re-run. This yielded more normally distributed residuals (sTable 1 and sFigure 1). Hence, we decided to use log-transformed $V_T$ values in the statistical analyses presented in the main article.

**sTable 1.** Shapiro-Wilk Normality test results for **(A)** untransformed data and **(B)** log transformed data.

| (A) | Parameter | W | *p* | (B) | Parameter | W | *p* |
|---|---|---|---|---|---|---|---|
| | **Grey matter $V_T$** | 0.941 | p<0.001 | | **log Grey matter $V_T$** | 0.985 | 0.116 |
| | **Frontal cortex $V_T$** | 0.940 | p<0.001 | | **log Frontal cortex $V_T$** | 0.983 | 0.085 |
| | **Temporal cortex $V_T$** | 0.944 | p<0.001 | | **log Temporal cortex $V_T$** | 0.987 | 0.211 |
| | **Occipital cortex $V_T$** | 0.934 | p<0.001 | | **log Occipital cortex $V_T$** | 0.983 | 0.088 |
| | **Parietal cortex $V_T$** | 0.937 | p<0.001 | | **log Parietal cortex $V_T$** | 0.984 | 0.106 |
| | **Hippocampus $V_T$** | 0.925 | p<0.001 | | **log Hippocampus $V_T$** | 0.986 | 0.152 |
| | **Thalamus $V_T$** | 0.938 | p<0.001 | | **log Thalamus $V_T$** | 0.985 | 0.13 |

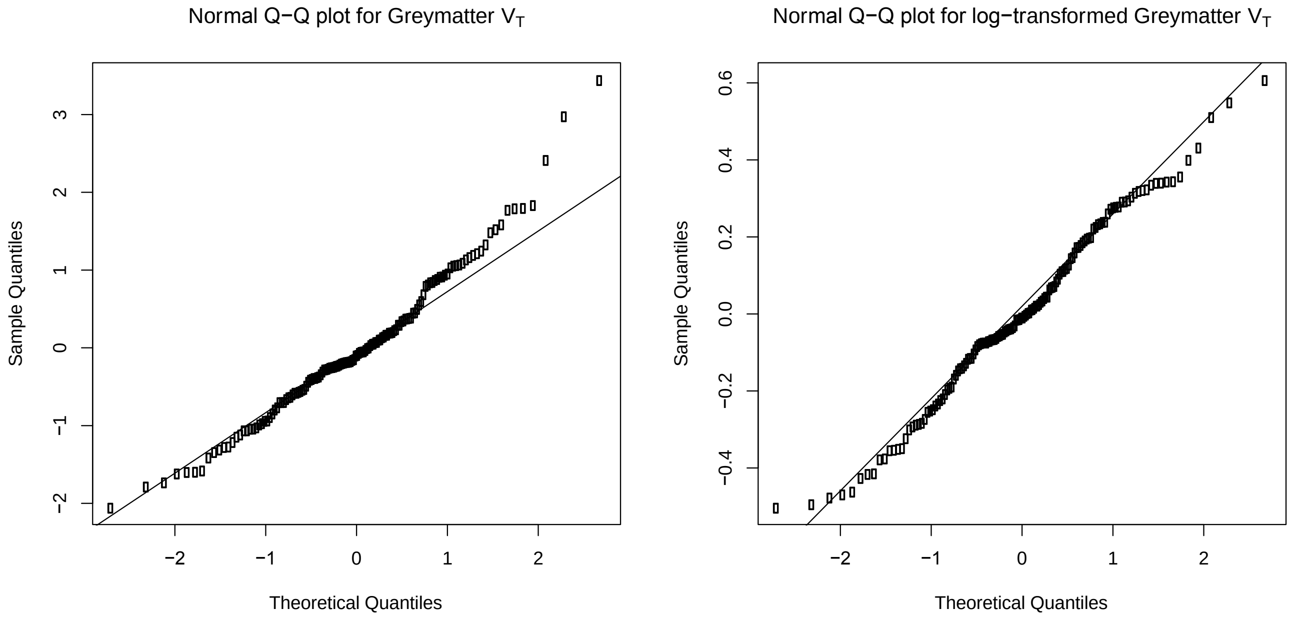


sFigure 1. QQ-plots of residuals from the linear-mixed-effects model using grey matter $V_T$ values (left) and log-transformed greymatter $V_T$ values (right).

*Confirmatory results*

For transparency, we also present the results from the main analysis using raw $V_T$ values below. These results should however be interpreted with caution, as the statistical models do not fulfill all assumptions (normally distributed residuals).

**sTable 2.** Fixed effects estimates with 95% confidence intervals and p-values for age, BMI and sex as regressors for regional [$^{11}$C]PBR28 raw untransformed $V_T$. N=140.

| Region | Age (β, [CI], p) | BMI (β, [CI], p) | Sex (β, [CI], p)* |
|---|---|---|---|
| **Grey matter** | 0.003, [-0.009,0.015], p=0.672 | -0.073, [-0.120,-0.026], p=0.003 | 0.641, [0.314,0.968], p<0.001 |
| **Frontal cortex** | 0.005, [-0.009,0.019], p=0.412 | -0.067, [-0.114,-0.020], p=0.007 | 0.668, [0.333,1.003], p<0.001 |
| **Temporal cortex** | 0.005, [-0.007,0.017], p=0.459 | -0.076, [-0.121,-0.031], p=0.001 | 0.615, [0.296,0.934], p<0.001 |
| **Occipital cortex** | 0.001, [-0.013,0.015], p=0.863 | -0.065, [-0.112,-0.018], p=0.009 | 0.687, [0.352,1.022], p<0.001 |
| **Parietal cortex** | 0.004, [-0.008,0.016], p=0.585 | -0.057, [-0.104,-0.010], p=0.018 | 0.684, [0.355,1.013], p<0.001 |
| **Hippocampus** | 0.003, [-0.011,0.017], p=0.639 | -0.093, [-0.150,-0.036], p=0.002 | 0.721, [0.313,1.129], p=0.001 |
| **Thalamus** | -0.002, [-0.014,0.01], p=0.734 | -0.085, [-0.136,-0.034], p=0.001 | 0.597, [0.238,0.956], p=0.001 |

*Coding: Males=0, Females=1

*Exploratory results*

sTable 3 presents the results of LME model for the age*sex interaction analysis. The interaction analysis showed a significant interaction between age and sex in all other regions apart occipital cortex and hippocampus.

**sTable 3.** Fixed effects estimates with 95% confidence intervals and p-values for age, BMI and age*sex as regressors for regional [$^{11}$C]PBR28 log-transformed $V_T$ in a separate subgroup analysis for males and females. N = 140.

| Region | Age (β, [CI], p) | BMI (β, [CI], p) | Sex (β, [CI], p)* | Age*Sex (β, [CI], p)* |
|---|---|---|---|---|
| **Grey matter** | 0.001, [-0.003,0.005], p=0.784 | -0.396, [-0.627,-0.165], p=0.001 | -0.022, [-0.034,-0.010], p<0.001 | -0.005, [-0.009,-0.001], p=0.031 |
| **Frontal cortex** | 0.001, [-0.003,0.005], p=0.571 | -0.416, [-0.651,-0.181], p=0.001 | -0.020, [-0.032,-0.008], p=0.001 | -0.005, [-0.009,-0.001], p=0.027 |
| **Temporal cortex** | 0.001, [-0.003,0.005], p=0.543 | -0.400, [-0.633,-0.167], p=0.001 | -0.023, [-0.035,-0.011], p<0.001 | -0.005, [-0.009,-0.001], p=0.028 |
| **Occipital cortex** | 0.000, [-0.004,0.004], p=0.937 | -0.373, [-0.600,-0.146], p=0.002 | -0.019, [-0.031,-0.007], p=0.001 | -0.004, [-0.008,0.000], p=0.054 |
| **Parietal cortex** | 0.000, [-0.004,0.004], p=0.877 | -0.431, [-0.660,-0.202], p<0.001 | -0.017, [-0.029,-0.005], p=0.004 | -0.005, [-0.009,-0.001], p=0.020 |
| **Hippocampus** | 0.000, [-0.004,0.004], p=0.891 | -0.334, [-0.585,-0.083], p=0.010 | -0.026, [-0.038,-0.014], p<0.001 | -0.004, [-0.008,0.000], p=0.108 |
| **Thalamus** | 0.001, [-0.003,0.005], p=0.782 | -0.409, [-0.660,-0.158], p=0.002 | -0.023, [-0.035,-0.011], p<0.001 | -0.005, [-0.009,-0.001], p=0.027 |

*Coding: Males=0, Females=1

*Robustness test*

To examine the robustness of our results, we performed the LME analysis also by excluding the subjects who had cholesterol medication (n=11), hormone replacement therapy (n=2) and who were current smokers (n=3) (sTable4; 16 excluded subjects, thus resulting total number of subjects n=124).

**sTable 4.** Fixed effects estimates with 95% confidence intervals and p-values for age, BMI and sex as regressors for regional [$^{11}$C]PBR28 log-transformed $V_T$, excluding smokers and subjects on medication. N=124.

| Region | Age (β, [CI], p) | BMI (β, [CI], p) | Sex (β, [CI], p)* |
|---|---|---|---|
| **Grey matter** | 0.003, [-0.001,0.007], p=0.078 | -0.023, [-0.035,-0.011], p<0.001 | 0.176, [0.088,0.264], p<0.001 |
| **Frontal cortex** | 0.004, [0.000,0.008], p=0.033 | -0.022, [-0.036,-0.008], p=0.001 | 0.185, [0.095,0.275], p<0.001 |
| **Temporal cortex** | 0.004, [0.000,0.008], p=0.031 | -0.024, [-0.036,-0.012], p<0.001 | 0.172, [0.082,0.262], p<0.001 |
| **Occipital cortex** | 0.002, [-0.002,0.006], p=0.159 | -0.020, [-0.032,-0.008], p=0.002 | 0.181, [0.095,0.267], p<0.001 |
| **Parietal cortex** | 0.003, [-0.001,0.007], p=0.081 | -0.019, [-0.031,-0.007], p=0.004 | 0.193, [0.105,0.281], p<0.001 |
| **Hippocampus** | 0.002, [-0.002,0.006], p=0.358 | -0.026, [-0.040,-0.012], p<0.001 | 0.161, [0.065,0.257], p=0.001 |
| **Thalamus** | 0.003, [-0.001,0.007], p=0.077 | -0.023, [-0.037,-0.009], p=0.001 | 0.161, [0.065,0.257], p=0.001 |

*Coding: Males=0, Females=1

sTable 5 presents results from the same robustness tests as described above but using raw untransformed $V_T$ values instead of log-transformed data.

**sTable 5.** Fixed effects estimates with 95% confidence intervals and p-values for age, BMI and sex as regressors for regional [$^{11}$C]PBR28 raw $V_T$, excluding smokers and subjects on medication. N=124.

| Region | Age (β, [CI], p) | BMI (β, [CI], p) | Sex (β, [CI], p)* |
|---|---|---|---|
| **Grey matter** | 0.004, [-0.010,0.018], p=0.598 | -0.076, [-0.127,-0.025], p=0.004 | 0.736, [0.375,1.097], p<0.001 |
| **Frontal cortex** | 0.006, [-0.008,0.020], p=0.355 | -0.071, [-0.124,-0.018], p=0.008 | 0.761, [0.394,1.128], p<0.001 |
| **Temporal cortex** | 0.005, [-0.009,0.019], p=0.414 | -0.078, [-0.127,-0.029], p=0.003 | 0.703, [0.350,1.056], p<0.001 |
| **Occipital cortex** | 0.002, [-0.012,0.016], p=0.775 | -0.068, [-0.121,-0.015], p=0.013 | 0.783, [0.413,1.153], p<0.001 |
| **Parietal cortex** | 0.005, [-0.009,0.019], p=0.509 | -0.062, [-0.113,-0.011], p=0.019 | 0.776, [0.413,1.139], p<0.001 |
| **Hippocampus** | 0.004, [-0.010,0.018], p=0.593 | -0.093, [-0.156,-0.030], p=0.005 | 0.836, [0.383,1.289], p<0.001 |
| **Thalamus** | -0.002, [-0.014,0.01], p=0.807 | -0.085, [-0.140,-0.030], p=0.003 | 0.693, [0.293,1.093], p=0.001 |

*Coding: Males=0, Females=1

sTable 6 presents the LME model for male and female subgroup analyses and the age*sex interaction analysis results. The subgroup analysis showed significant positive associations between age and log-transformed $V_T$ in all regions apart from hippocampus in male subjects, whereas for female subjects no significant associations were found. Furthermore, significant negative association between BMI and log-transformed $V_T$ was observed in males in all regions, but for females the association was significant only in hippocampus. Despite the sex-related differences in the subgroup analysis of age effect, the interaction analysis didn't show a significant interaction between age and sex.

**sTable 6.** Fixed effects estimates with confidence intervals and p-values for age and BMI as regressors for regional [11C]PBR28 log-transformed $V_T$ in a separate subgroup analysis for males and females (**A**). Fixed effects estimates with 95% confidence intervals and p-values in a separate subgroup analyses for males (N = 67) and females (N = 57) for regional [11C]PBR28 log-transformed $V_T$. (**B**). Fixed effect estimates for age, BMI and an age*sex interaction effect. N=124.

| (A) | Region | Age (β, [CI], p), Sex = M | Age (β, [CI], p), Sex = F | BMI (β, [CI], p), Sex = M | BMI (β, [CI], p), Sex = F |
|---|---|---|---|---|---|
| | **Grey matter** | 0.006, [0.002,0.010], p=0.003 | -0.002, [-0.008,0.004], p=0.586 | -0.025, [-0.041,-0.009], p=0.003 | -0.018, [-0.038,0.002], p=0.073 |
| | **Frontal cortex** | 0.007, [0.003,0.011], p=0.001 | -0.001, [-0.007,0.005], p=0.696 | -0.023, [-0.039,-0.007], p=0.007 | -0.018, [-0.038,0.002], p=0.084 |
| | **Temporal cortex** | 0.007, [0.003,0.011], p=0.001 | -0.001, [-0.007,0.005], p=0.763 | -0.025, [-0.041,-0.009], p=0.003 | -0.020, [-0.040,0.000], p=0.054 |
| | **Occipital cortex** | 0.005, [0.001,0.009], p=0.008 | -0.002, [-0.008,0.004], p=0.514 | -0.022, [-0.038,-0.006], p=0.006 | -0.015, [-0.035,0.005], p=0.137 |
| | **Parietal cortex** | 0.006, [0.002,0.010], p=0.003 | -0.002, [-0.008,0.004], p=0.523 | -0.021, [-0.037,-0.005], p=0.015 | -0.014, [-0.034,0.006], p=0.164 |
| | **Hippocampus** | 0.004, [0.000,0.008], p=0.051 | -0.002, [-0.006,0.002], p=0.433 | -0.027, [-0.045,-0.009], p=0.006 | -0.020, [-0.038,-0.002], p=0.040 |
| | **Thalamus** | 0.005, [0.001,0.009], p=0.009 | -0.001, [-0.007,0.005], p=0.757 | -0.022, [-0.040,-0.004], p=0.015 | -0.021, [-0.041,-0.001], p=0.057 |

| (B) | Region | Age (β, [CI], p) | BMI (β, [CI], p) | Sex (β, [CI], p)* | Age*Sex (β, [CI], p)* |
|---|---|---|---|---|---|
| | **Grey matter** | 0.001, [-0.003,0.005], p=0.661 | -0.378, [-0.621,-0.135], p=0.003 | -0.023, [-0.035,-0.011], p=0.001 | -0.004, [-0.008,0.000], p=0.085 |
| | **Frontal cortex** | 0.002, [-0.002,0.006], p=0.475 | -0.398, [-0.647,-0.149], p=0.002 | -0.022, [-0.034,-0.010], p=0.001 | -0.004, [-0.008,0.000], p=0.074 |
| | **Temporal cortex** | 0.001, [-0.003,0.005], p=0.474 | -0.385, [-0.632,-0.138], p=0.003 | -0.024, [-0.036,-0.012], p<0.001 | -0.004, [-0.008,0.000], p=0.072 |
| | **Occipital cortex** | 0.001, [-0.003,0.005], p=0.790 | -0.355, [-0.596,-0.114], p=0.005 | -0.020, [-0.032,-0.008], p=0.002 | -0.003, [-0.007,0.001], p=0.133 |
| | **Parietal cortex** | 0.001, [-0.003,0.005], p=0.748 | -0.415, [-0.658,-0.172], p=0.001 | -0.018, [-0.030,-0.006], p=0.005 | -0.004, [-0.008,0.000], p=0.058 |
| | **Hippocampus** | 0.000, [-0.004,0.004], p=0.983 | -0.314, [-0.579,-0.049], p=0.022 | -0.026, [-0.040,-0.012], p<0.001 | -0.003, [-0.007,0.001], p=0.230 |
| | **Thalamus** | 0.001, [-0.003,0.005], p=0.718 | -0.393, [-0.662,-0.124], p=0.005 | -0.023, [-0.037,-0.009], p=0.001 | -0.005, [-0.011,0.001], p=0.072 |

*Coding: Males=0, Females=1

sTable 7 presents the GM results of LME model, where centre was inserted as a dummy predictor and as an interaction-effect to age, BMI and sex. The age interaction estimate for Turku data was statistically significant, whereas the other interaction estimates were not significant

**sTable 7.** (**A**) Fixed effects estimates with 95% confidence intervals and p-values for age for age, BMI, sex, centre and age*centre as regressors for gray matter [$^{11}$C]PBR28 log-transformed $V_T$. (**B**) Fixed effects estimates with 95% confidence intervals and p-values for age for age, BMI, sex, centre and BMI*centre as regressors for gray matter [$^{11}$C]PBR28 log-transformed $V_T$. (**C**) Fixed effects estimates with 95% confidence intervals and p-values for age for age, BMI, sex, centre and sex*centre as regressors for gray matter [$^{11}$C]PBR28 log-transformed $V_T$. N=140. All variables were standardized before the interaction analysis.

| (A) | Effect | β, [CI], *p* |
|---|---|---|
| | **Age** | 0.225, [0.045,0.405], p=0.016 |
| | **BMI** | 0.400, [0.200,0.600], p<0.001 |
| | **Sex** | -0.221, [-0.331,-0.111], p<0.001 |
| | **Center=Turku** | 0.589, [0.201,0.977], p=0.004 |
| | **Center=Yale** | 1.031, [0.525,1.537], p<0.001 |
| | **Age*Center=Turku** | -0.536, [-0.955,-0.117], p=0.013 |
| | **Age*Center=Yale** | 0.054, [-0.367,0.475], p=0.804 |

| (B) | Effect | β, [CI], *p* |
|---|---|---|
| | **Age** | 0.157, [-0.006,0.32], p=0.061 |
| | **BMI** | 0.381, [0.171,0.591], p=0.001 |
| | **Sex** | -0.257, [-0.494,-0.02], p=0.036 |
| | **Center=Turku** | 0.263, [-0.039,0.565], p=0.090 |
| | **Center=Yale** | 0.923, [0.576,1.27], p<0.001 |
| | **Age*Center=Turku** | 0.018, [-0.260,0.296], p=0.900 |
| | **Age*Center=Yale** | 0.103, [-0.215,0.421], p=0.527 |

| (C) | Effect | β, [CI], *p* |
|---|---|---|
| | **Age** | 0.151, [-0.006,0.308], p=0.061 |
| | **BMI** | 0.384, [0.051,0.717], p=0.026 |
| | **Sex** | -0.225, [-0.337,-0.113], p<0.001 |
| | **Center=Turku** | 0.300, [-0.055,0.655], p=0.100 |
| | **Center=Yale** | 0.845, [0.457,1.233], p<0.001 |
| | **Age*Center=Turku** | -0.083, [-0.53,0.364], p=0.716 |
| | **Age*Center=Yale** | 0.257, [-0.333,0.847], p=0.395 |

*Coding: Males=0, Females=1

### Collaborators

Members of the HRRT [$^{11}$C]PBR28 study group:

Jouni Tuisku[1], Pontus Plavén-Sigray[2], Edward C. Gaiser[4,5], Laura Airas[1,6], Haidar Al-Abdulrasul[1], Anna Brück[1,6], Richard E. Carson[4], Ming-Kai Chen[4,5], Karin Collste[2], Kelly P. Cosgrove[4,5], Laura Ekblad[1], Irina Esterlis[4,5], Lars Farde[2,7], Anton Forsberg[2], Jean-Dominique Gallezot[4], Christer Halldin[2], Semi Helin[1], Ansel Hillmer[4,5], Yiyun Huang[4], Caroline O. Höglund[2,9,10], Jarkko Johansson[1], Aurelija Jucaite[2,7], Eva Kosek[8], Jon Lampa[9], Mats Lekander[3,8], Noora Lindgren[1], Päivi Marjamäki[1], Nabeel Nabulsi[4], Brian Pittman[5], Eero Rissanen[1,6], Christine M. Sandiego[4,5], Per Stenkrona[2,] Marcus Sucksdorff[1,6], Andrea Varrone[2,3], Juha Rinne[1,6], David Matuskey[4,5], Simon Cervenka[2]

[1]*Turku PET Centre, University of Turku, Turku, Finland.*

[2]*Centre for Psychiatry Research, Department of Clinical Neuroscience, Karolinska Institutet and Stockholm County, Stockholm, Sweden.*

[3]*Stress Research Institute, Department of Psychology, Stockholm University*

[4]*PET Center, Department of Radiology and Biomedical Imaging, Yale University, New Haven, CT 06520.*
[5]*Department of Psychiatry, Yale University, New Haven, CT 06511.*
[6]*Division of Clinical Neurosciences, Turku University Hospital, Turku, Finland*
[7]*PET Science Centre, Precision Medicine and Genomics, IMED Biotech Unit, AstraZeneca, Karolinska Institutet, Sweden*
[8]*Department of Clinical Neuroscience, Karolinska Institutet, Stockholm, Sweden*
[9] *Department of Medicine and Center for Molecular Medicine, Karolinska Institutet, Karolinska University Hospital, Stockholm, Sweden*
[10]*Department of Physiology and Pharmacology, Karolinska Institutet, Stockholm, Sweden*
